# Insulin-like peptide receptor mediated signaling pathways orchestrate regulation of energy homeostasis in the Pacific oyster, *Crassostrea gigas*

**DOI:** 10.1101/2021.02.25.432956

**Authors:** Yongjing Li, Huiru Fu, Fuqiang Zhang, Liting Ren, Jing Tian, Qi Li, Shikai Liu

**Affiliations:** Key Laboratory of Mariculture (Ocean University of China), Ministry of Education, and College of Fisheries, Ocean University of China, Qingdao 266003, China; Laboratory for Marine Fisheries Science and Food Production Processes, Qingdao National Laboratory for Marine Science and Technology, Qingdao 266237, China

**Keywords:** *Crassostrea gigas*, IIS, nutrient abundance, temperature, energy metabolism

## Abstract

The involvement of insulin/insulin-like growth factor (IIS) signaling pathway in growth regulation of marine invertebrates remains largely unexplored. In this study, we used a fast-growing Pacific oyster (*Crassostrea gigas*) variety “Haida No.1” as material to unravel the role of IIS system in growth regulation in oysters. Systematic bioinformatics analyses allowed to identify major components of IIS signaling pathway and insulin-like peptide receptor (ILPR) mediated signaling pathways, including PI3K-AKT, RAS-MAPK, and TOR, in *C. gigas*. Expression levels of the major genes in IIS and its downstream signaling pathways were significantly higher in “Haida No.1” than wild oysters, suggesting their involvement in growth regulation of *C. gigas*. Expression profiles of IIS and its downstream signaling pathway genes were significantly altered by nutrient abundance and culture temperature. These results suggested that IIS signaling pathway coupled with the ILPR mediated signaling pathways orchestrated energy homeostasis to regulate growth in the Pacific oyster.

**Research Highlights:**

1. *ILPR, IRS, IGFBPRP,* and *IGFALS* genes were characterized in the *C. gigas*.
2. Major genes of IIS signaling pathway were highly expressed in fast-growing *C. gigas*.
3. IIS and downstream pathways participates in energy homeostasis of oysters.
4. ILPR mediated signaling pathways orchestrate growth regulation in oysters.

## 1. Introduction

Nutrient abundance is one of the most important environmental factors that are critical to growth and reproduction of all organisms. Nutrient sensing and energy metabolism are coordinated by networks of signaling cascades such as insulin/insulin-like growth factor signaling (IIS), target of rapamycin (TOR), and adenosine monophosphate-activated protein kinase (AMPK) signaling pathways [1]. The IIS and AMPK signaling pathways are two critical coordinators of energy homeostasis and metabolic processes across diverse vertebrate and invertebrate species. The TOR signaling pathway integrates various environmental factors and the signal transducing from the IIS and AMPK pathways to direct the cell growth [1, 2].

As one of the most important nutrient sensors, the mechanism of IIS pathway in growth regulation of vertebrates has been well studied. The IIS signaling pathway, including insulin/insulin-like growth factors (IGFs), insulin/IGF receptor (INSR, ILPR, IGFR), insulin receptor substrate (IRS), insulin-like growth factor-binding proteins (IGFBPs), and insulin-like growth factor-binding protein complex acid labile subunits (IGFALSs), regulates multiple cellular processes, including cell growth and proliferation, energy metabolism, hormone secretion, and glucose homeostasis [3–5]. In vertebrates, the insulin acutely alters in response to the nutrient abundance and other circulating factors to regulate the nutrient uptake and storage activity of organisms [6]. The IGFs are regulated by the growth hormone (GH) which is secreted by neuroendocrine cells of the anterior pituitary gland and function to regulate the growth of organisms [7, 8]. The levels of IGFs in serum are strictly controlled by the ternary complex “IGF-IGFBP-ALS” [9]. Binding of insulin or IGFs to the tyrosine kinase receptor (INSR, IGFR) leads to activation of receptor and phosphorylation of cellular receptor substrate (IRS) which cooperate to trigger the PI3K-AKT and RAS-MAPK pathways to regulate the cell proliferation, glycogen metabolism and protein synthesis [4, 10].

Only insulin-like peptide but no growth hormone has been identified in various invertebrates [11–13]. The insulin-like peptides were produced from brain or small clusters of neuroendocrine cells, and involved in the neuroendocrine activity in several insects and gastropod mollusks [14–18]. The insulin-like peptide receptors and various members of IGFBP in invertebrates, which share similar structure and function with their counterparts in vertebrates, have also been identified [19–27]. Furthermore, genes involved in PI3K-AKT and RAS-MAPK signaling pathways, such as *PI3K, AKT, PTEN,* and *RAS* were also identified in invertebrates [28–30]. Levels of expression or phosphorylation of key components of these pathways has been associated with growth difference [29, 31–35]. A recent study in pearl oyster reported that the insulin-like peptide recombinant protein induced the expressions of *ILPR* and the major genes of PI3K-AKT and RAS-MAPK pathways, suggesting a functional IIS signaling cascade mediated by *ILPR* [36]. Therefore, systemic characterization of IIS signaling and its downstream pathways that are involved in growth would be essential for unravelling molecular mechanisms of growth regulation in invertebrates such as mollusks.

The Pacific oyster (*Crassostrea gigas*) is one of the most widely cultured marine mollusks, which has been introduced from Asia to many other countries around the world [37]. A selection breeding program of the *C. gigas* in China performed for over ten years have produced a fast-growing variety named as “Haida No.1”, which exhibits significant growth advantages over wild oysters [38, 39]. The fast-growing variety provides an ideal research system for growth studies in oyster. In a previous work, we identified four insulin-like peptide genes in *C. gigas,* and showed that expression levels of ILPs were significantly higher in fast-growing “Haida No.1” than wild oysters [40]. Environmental factors such as nutrient abundance and ambient temperature had significant effects on expression of ILP genes and growth of oysters. To further understand the roles of IIS signaling pathway in growth regulation of oysters, in this work, we performed an extensive multi-omics data mining to identify key components of IIS, PI3K-AKT and RAS-MAPK signaling pathways in *C. gigas*. Expression profiles of major genes of the IIS and ILPR mediated signaling pathways were determined in “Haida No.1” and wild oysters. Fasting/re-feeding and low temperature culture experiments were performed to further understand how the IIS signaling pathway was altered by nutrient level and temperature to affect growth of *C. gigas*. This work provided a systemic characterization of IIS and ILPR mediated signaling pathways involved in growth of *C. gigas*, which would provide valuable information toward understanding of molecular basis of growth regulation in oysters and other invertebrates.

## 2. Materials and methods

### 2.1 Sequence analysis

The amino acid sequences of IIS signaling pathway genes, including *ILPR, IRS, IGFBP* and *IGFALS,* from *Homo sapiens, Gallus gallus, Xenopus laevis, Anolis carolinensis, Danio rerio, Limulus polyphemus, Musca domestica, Tetranychus urticae, Aplysia californica, Mizuhopecten yessoensis, Octopus bimaculoid, Pomacea canaliculata, Crassostrea virginica* and *Crassostrea gigas* were retrieved from the NCBI database. The detailed information of these sequences was provided in Supplementary Table 1. The sequences were aligned using ClustalW2 (http://www.ebi.ac.uk/Tools/msa/clustalw2/), and the phylogenetic trees were constructed using neighbor-joining (NJ) approach in MEGA7 [41]. The reliability of topological structure was tested using 1000 bootstrap replications.

### 2.2 Real-time PCR and statistical analysis

All the primers used for real-time PCR were designed by Primer Express software (Applied Biosystems, USA) and provided in Supplementary Table 2. The 2-fold dilutions of cDNA isolated from adductor muscle and the internal control gene *elongation factor 1α (EF1α)* were used to assess primer efficiency and the primer sets with an efficiency of 90-110% were used for real-time PCR analysis. The real-time PCR reactions were carried out in a LightCycler 480 machine (Roche, Switzerland) in a 20 μL system with a mixture of 10 μL 2×SYBR Premix ExTaq (Qiagen, Germany), 2.0 μL diluted cDNA or double distilled water as a negative control, 6 μL PCR-grade water, and 1.0 μL each primer (10 μM). The PCR reactions were initiated by denaturation at 95 °C for 3 min, followed by 40 amplification cycles at 95 °C for 15 s and 60 °C for 30 s. Dissociation protocols were used to measure melting curves. The relative expression level (RNA abundance) was calculated by dividing the copy number of the target gene by that of the internal control gene. Data were expressed as the mean ± SD. Significant differences (*P* < 0.05) were determined using the one-way ANOVA and Student’s *t-*test for single or multiple comparisons, respectively.

### 2.3. Expression profiles of the major genes in IIS, PI3K-AKT and RAS-MAPK signaling pathways

Expression profiles of the major genes in IIS, PI3K-AKT and RAS-MAPK signaling pathways, including *ILPR, IRS, IGFBPRP, IGFALS, PI3K, PDK, AKT, RAS, MEK, ERK, PTEN, TOR, FoxO, GSK3β,* and *S6K,* were determined in the adductor muscle of “Haida No.1” and wild oysters. Total RNA was extracted using Trizol Reagent (Invitrogen) and reversely transcribed into cDNA according to the PrimeScript™ RT reagent Kit with gDNA Eraser (Perfect Real Time) (TaKaRa, Japan) manufacturer’s instructions. Real-time PCR was carried out as described above.

### 2.4 Fasting and re-feeding culture experiment

Ninety 8-mounth-old *C. gigas* were randomly divided into three groups and used for fasting and re-feeding experiment. Six oysters collected before the experiment were used as control (as indicated by C), the other oysters were starved for 14 days and then refed with frozen *Chlorella* ad libitum for 36 hours. Samples were collected on day 1, 3, 5, 7 and 14 (as indicated by F1d, F3d, F5d, F7d and F14d, respectively) during fasting process, and collected at 1, 3, 6, 12, 24 and 36 hours (as indicated by R1h, R3h, R6h, R12h, R24h, R36h, respectively) after re-feeding with six oysters at each time point. Various tissues including the labial palp, gill, mantle, digestive gland, hematocyte, heart, visceral ganglia, and adductor muscle were rapidly excised, frozen in liquid nitrogen, and then stored at −80 °C until use. Total RNA extraction and cDNA synthesis were performed as described above. The cDNA from the all eight tissues at each sampling point was pooled into one sample with equal amount per tissue, and used for real-time PCR analysis according to the method described above.

### 2.5 Low temperature culture experiment

Oyster umbo larvae with similar size (2.5±0.3 mm) were stocked into three 75 L cylindrical polyethylene vessels (larval density of 4/mL) and reared at 5 ± 1 °C, 15 ± 1 °C and 25 ±1 °C, respectively. Light was supplied by fluorescent lamps with 12:12 Light: Dark photoperiod. For each culture temperature, three random replicates were allocated, the survival and feeding situation of the larvae was monitored daily for 10-day culture, the shell height of 30 randomly collected larvae was measured, and the whole larvae were collected and deposited into RNAstore (CWBIO, China) until used for RNA expression analyses.

## 3. Results

### 3.1 Identification of ILPR, IRS, IGFBP, and IGFALS in C. gigas

One *ILPR,* one *IRS*, one *IGFBP,* and seven *IGFALS* genes were identified in *C. gigas* through extensive genome and transcriptome data mining. The gene names, sequence characteristics and accessions were provided in Table 1. Phylogenetic analysis showed that the *C. gigas* ILPR was clustered into one clade with the ILPRs of arthropods and other mollusks, the vertebrates IGF1R, IGF2R, INSR, and INSRR were clustered into separate clades, respectively (Fig. 1). In addition, the IRS of *C. gigas* was clustered into one clade with the IRS of arthropods and other mollusks, while the three IRSs in vertebrates, IRS1, IRS2 and IRS4 were clustered into other separate clades, respectively (Supplementary Fig. 1). The only one IGFBPRP in *C. gigas* was clustered into the clade with vertebrate IGFBP7 (also known as IGFBPRP1) and IGFBPRP from other mollusks, while the other six vertebrate-specific IGFBP members were clustered into other separate clades, respectively (Supplementary Fig. 2).

**Fig. 1.**
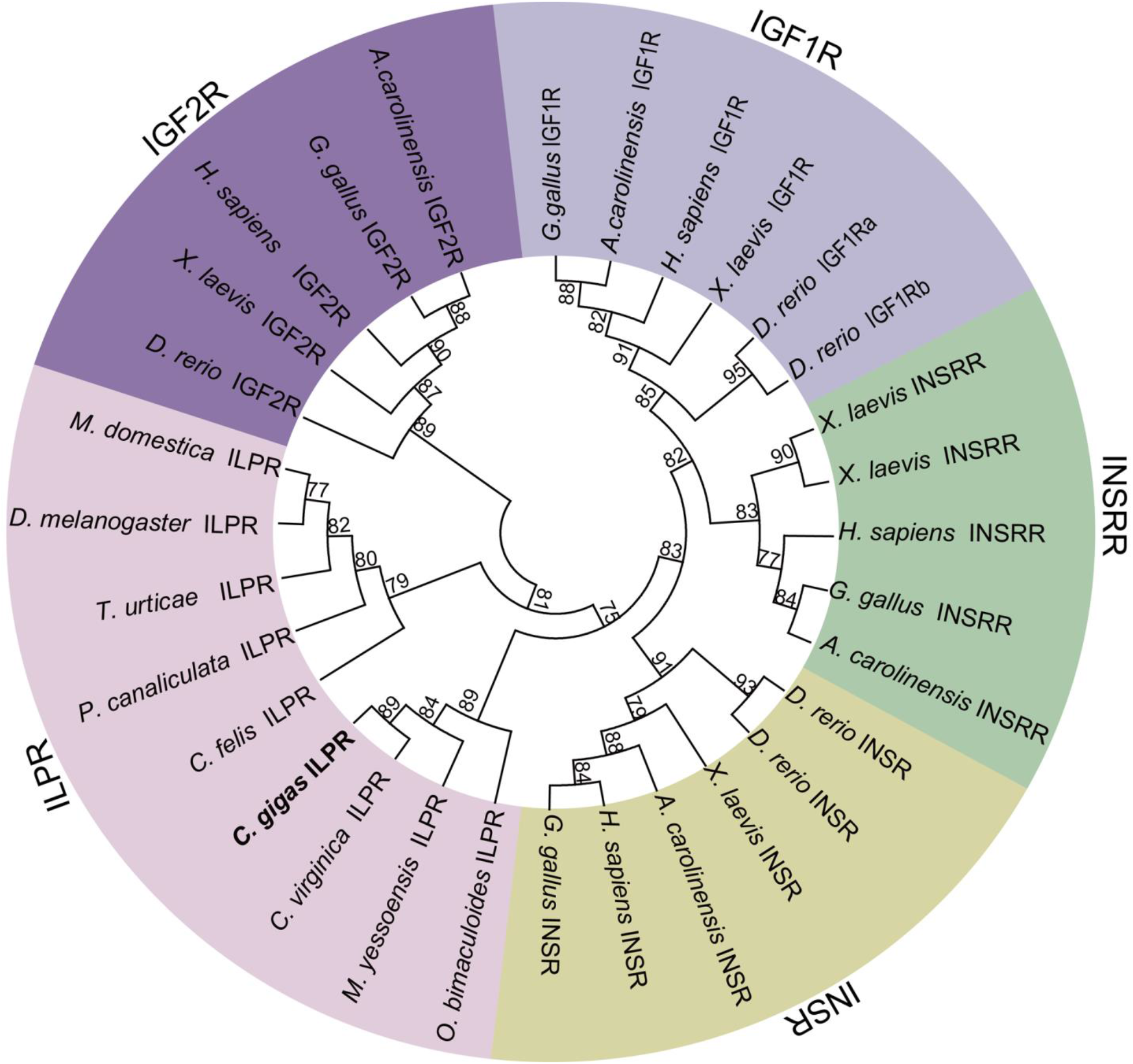
Phylogenetic analyses of the insulin receptors and insulin-like peptide receptors. The amino acid sequences of insulin receptors and insulin-like peptide receptors from *Homo sapiens, Gallus gallus, Xenopus laevis, Anolis carolinensis, Danio rerio, Limulus polyphemus, Musca domestica, Ctenocephalides felis, Mizuhopecten yessoensis, Tetranychus urticae, Octopus bimaculoid*, *Pomacea canaliculata*, *Crassostrea virginica* and *Crassostrea gigas* were retrieved from the NCBI database. Phylogenetic tree was constructed using neighbor-joining (NJ) approach in MEGA7. The reliability of topological structure was tested using 1000 bootstrap replications.

**Table 1.** Sequence characteristics of the IIS signaling pathway genes in *Crassostrea gigas*

| Gene name | Gene full name | Amino acid (aa) | mRNA (bp) | ORF (bp) | NCBI Gene ID |
| --- | --- | --- | --- | --- | --- |
| *ILPR* | Insulin-like peptide receptor | 1561 | 9092 | 4686 | 105348544 |
| *IRS* | Insulin receptor substrate 1-B | 1400 | 6375 | 4203 | 105342662 |
| *IGFBPRP* | Insulin-like growth factor-binding protein-related protein 1 | 258 | 1926 | 777 | 105339347 |
| *ALS5594* | Insulin-like growth factor-binding protein complex acid labile subunit | 978 | 4175 | 2937 | 105319673 |
| *ALS7789* | Insulin-like growth factor-binding protein complex acid labile subunit | 780 | 2607 | 2343 | 105342519 |
| *ALS7489* | Insulin-like growth factor-binding protein complex acid labile subunit | 1000 | 3987 | 3003 | 105342288 |
| *ALS2466* | Insulin-like growth factor-binding protein complex acid labile subunit | 913 | 4713 | 2744 | 105338877 |
| *ALS6223* | Insulin-like growth factor-binding protein complex acid labile subunit | 349 | 1726 | 1050 | 105320119 |
| *ALS1089* | Insulin-like growth factor-binding protein complex acid labile subunit | 456 | 4620 | 1371 | 105324304 |
| *ALS4336* | Insulin-like growth factor-binding protein complex acid labile subunit | 858 | 3189 | 2577 | 105331921 |

In striking contrast to the identification of only one IGFBP, seven IGFALSs were found in *C. gigas*. Phylogenetic analysis showed that the *C. gigas* ALS4336 was clustered into one clade with ALSs of *Mizuhopecten yessoensis,* the ALS7489 and ALS5594 in *C. gigas* were clustered into one clade with ALSs from the *Crassostrea virginica*, *Mizuhopecten yessoensis*, and *Octopus bimaculoid,* then the three ALSs in *C. gigas* were clustered into a larger clade with the ALSs from several vertebrates. Furthermore, the ALS1089, ALS2466, ALS7789 and ALS6223 in *C. gigas* were clustered into one clade with the ALS of *Crassostrea virginica* and had relatively closer homology to the ALS in invertebrates. The phylogenetic analysis also indicated that the ALS1089, ALS2466, ALS7789 and ALS6223 in *C. gigas* had a relatively primitive evolution status in comparison with other three ALSs, including the ALS4336, ALS7489 and ALS5594 (Supplementary Fig. 3).

### 3.2 Expression profiles of *IIS, PI3K-AKT and RAS-MAPK signaling pathway genes between “Haida No.1” and wild oysters*

The IIS signaling pathway genes including *ILPR*, *IGFBPRP* and *IRS* were all expressed at higher levels in fast-growing “Haida No. 1” than wild oysters (Fig. 2A). Among the seven *ALS* genes, the *ALS7789* and *ALS7489* were highly expressed in “Haida No.1”, while *ALS5594* was expressed at a higher level in wild oysters, and the other *ALS*s showed no expression difference between “Haida No.1” and wild oysters (Fig. 2B).

**Fig. 2.**
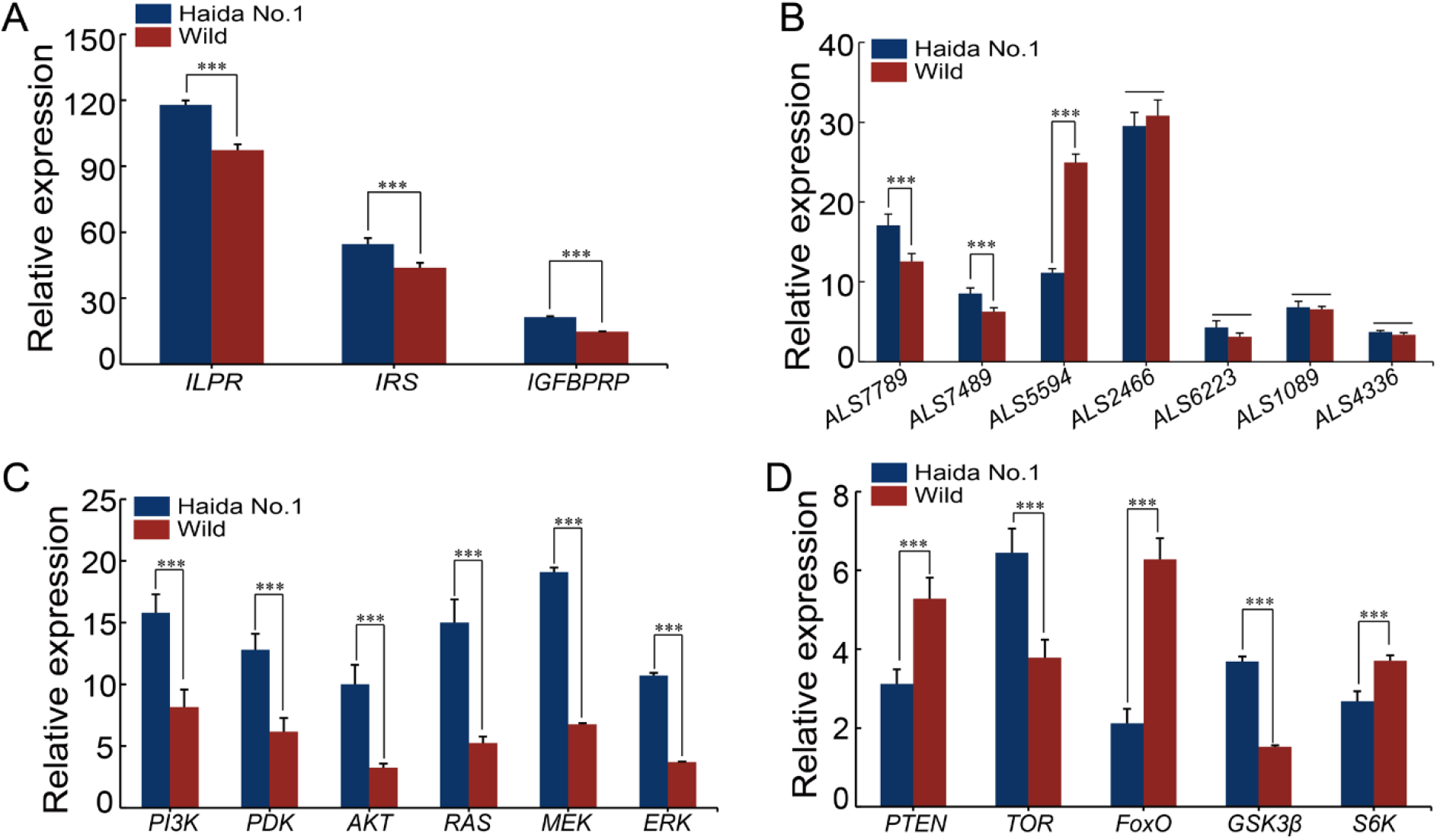
Expression profiles of the IIS, PI3K-AKT, RAS-MAPK, and TOR signaling pathway genes. Data are expressed as the mean ± SD (n = 4). The significant difference (*P* < 0.05) among groups is indicated by the asterisk.

The genes involved in PI3K-AKT and RAS-MAPK signaling pathways, including *PI3K, PDK, AKT, RAS, MEK,* and *ERK,* were all highly expressed in “Haida No.1” variety (Fig. 2C). Furthermore, downstream genes of the PI3K-AKT signaling pathway, including *GSK3β, FoxO, PTEN, TOR*, and *S6K,* showed distinct expression profiles between the “Haida No.1” and wild oysters, with *GSK3β* and *TOR* being expressed at higher levels in “Haida No.1”, while *PTEN, FoxO* and *S6K* being expressed at higher levels in the wild oysters (Fig. 2D).

### 3.3 Effect of nutrient abundance on the IIS, PI3K-AKT and RAS-MAPK signaling pathway genes in C. gigas

Expression profiles of *ILPR, IRS* and *IGFBPRP* genes were quite distinct in the fasting and re-feeding culture experiment. Specifically, *ILPR, IRS* and *IGFBP* were significantly down-regulated from day one to day 14 during the fasting treatment. Once refeeding, the expression of *ILPR* was significantly up-regulated in one hour and peaked at 12h and 24h, then decreased to a relative lower level at 36h (Fig. 3A), the expression of *IRS* was significantly up-regulated at 3h and peaked at 36h (Fig. 3B), the expression of *IGFBPRP* was significantly up-regulated in one hour and peaked at 3h, then decreased and maintained at a regular level comparable to that of control at 36h (Fig. 3C).

**Fig. 3.**
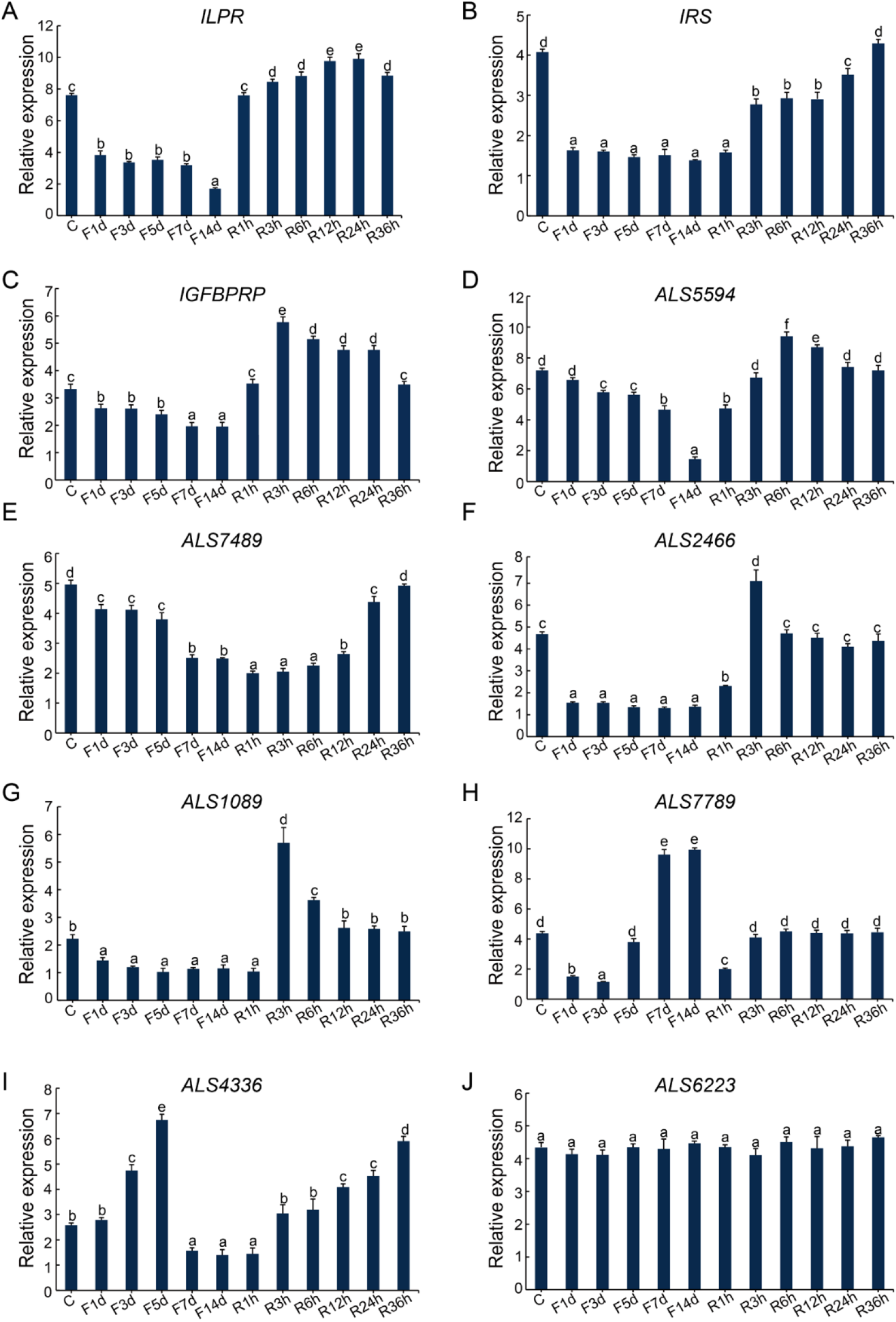
Effects of fasting and re-feeding on expressions of the IIS signaling pathway genes. C represented the control group, F1d, F3d, F5d, F7d and F14d represented 1, 3, 5, 7 and 14 days after fasting, respectively; R1h, R3h, R6h, R12h, R24h and R36h represented 1, 3, 6, 12, 24 and 36 hours after re-feeding, respectively. Data are expressed as the mean ± SD (n = 3). The significant difference (*P* < 0.05) among timepoints is indicated by the different lowercase letters.

The expressions of the seven *IGFALS*s showed gene-specific expression patterns after fasting and re-feeding treatment. Both *ALS5594, ALS7489, ALS2466* and *ALS1089* genes were all significantly down-regulated from day one to day 14 during fasting treatment and up-regulated in 1, 3 or 12 hours once re-feeding (Fig. 3D, E, F, G). In addition, the expression of *ALS7789* was firstly down-regulated from day one to day 3, and then up-regulated from day 5 to day 14 during the fasting process. Once re-feeding, its expression was firstly down-regulated, then up-regulated during 1 to 3 hours, and finally maintained at a control level (Fig. 3H). The expression of *ALS4366* was significantly up-regulated on day 3 and peaked on day 5, then decreased from day 7 to day 14 during the fasting process. Once refeeding, expression of *ALS4366* was up-regulated at 3h and peaked at 36h (Fig. 3I). The expression of *ALS6223* was not affected by fasting and re-feeding treatment (Fig. 3J).

The expression profiles of *TOR, S6K, FoxO* and *GSK3β* genes were quite distinct after the fasting and re-feeding treatment. The *TOR, S6K* and *GSK3β* showed stepwise decrease from day one to day 14 during fasting. Once refeeding, those genes were significantly up-regulated in one hour and peaked at 36h (Fig. 4A, B, C), the expression of *FoxO* was up-regulated continuously from day 1 to day 14 during fasting, then down-regulated from 3 h to 36 h to the control level once re-feeding (Fig. 4D).

**Fig. 4.**
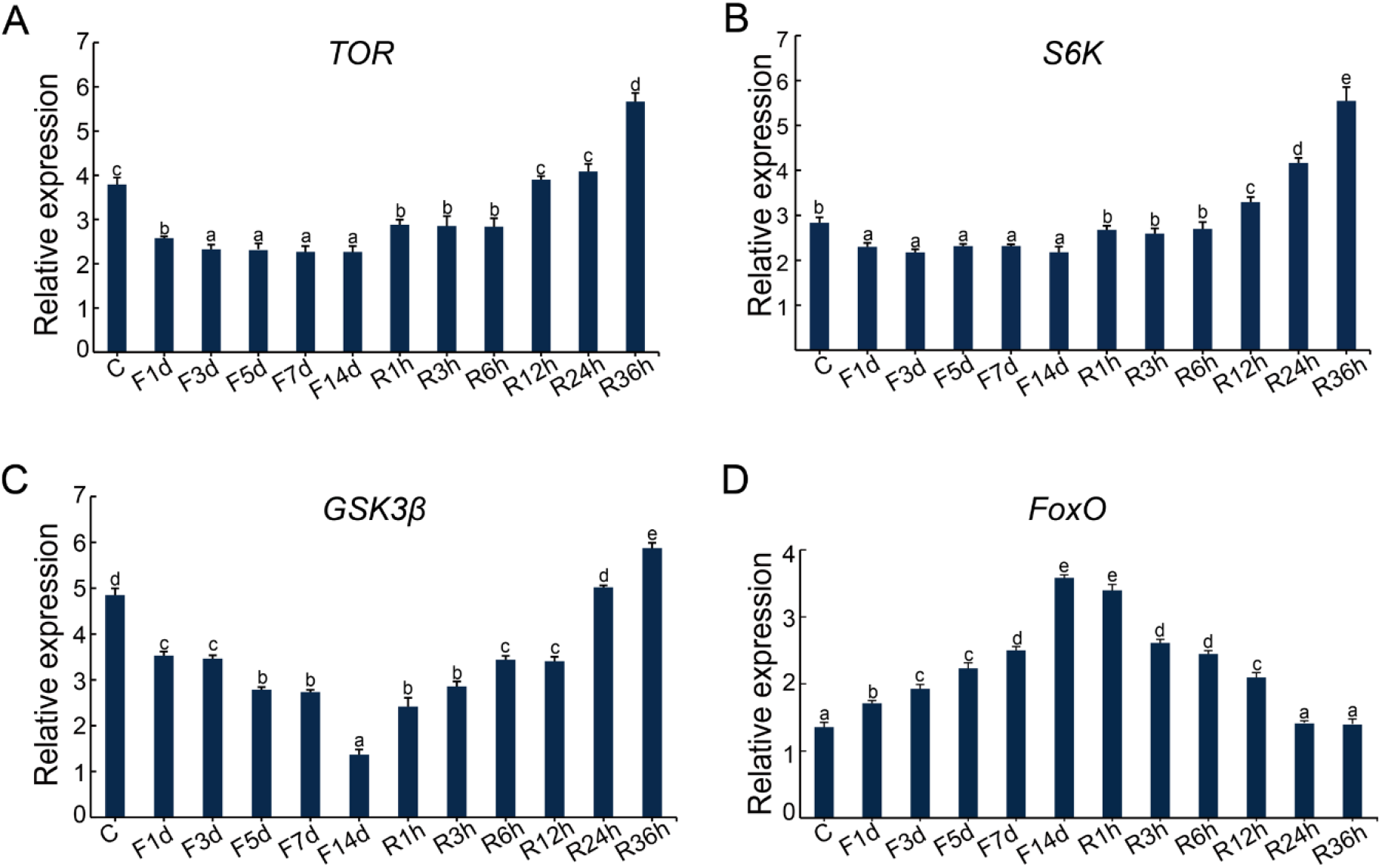
Effects of fasting and re-feeding on expressions of genes in the PI3K-AKT, RAS-MAPK, and TOR signaling pathways. C represented the control group, F1d, F3d, F5d, F7d and F14d represented 1, 3, 5, 7 and 14 days after fasting, respectively; R1h, R3h, R6h, R12h, R24h and R36h represented 1, 3, 6, 12, 24 and 36 hours after re-feeding, respectively. Data are expressed as the mean ± SD (n = 3). The significant difference (*P* < 0.05) among timepoints is indicated by the different lowercase letters.

### 3.4 Effect of temperature on the IIS, PI3K-AKT and RAS-MAPK signaling pathway genes in C. gigas

Low temperature significantly suppressed the growth of oyster larvae as well as the expression of the four insulin-like peptide genes [40]. Similarly, we found that the expression of the major IIS, PI3K-AKT and RAS-MAPK signaling pathway genes, including *ILPR, IRS, IGFBPRP, ALS7789, ALS7489, ALS1089, PI3K, PDK, AKT, RAS, MEK, ERK, PTEN, TOR, GSK3β* and *S6K* genes were also suppressed under the low culture temperature (Fig. 5A, B, C). In contrast, expression levels of *ALS5594*, *ALS2466*, *ALS6223* and *ALS4336* genes were not affected (Fig. 5B), while the *FoxO* was expressed at a relatively higher level under low temperature than normal temperature (Fig. 5D).

**Fig. 5.**
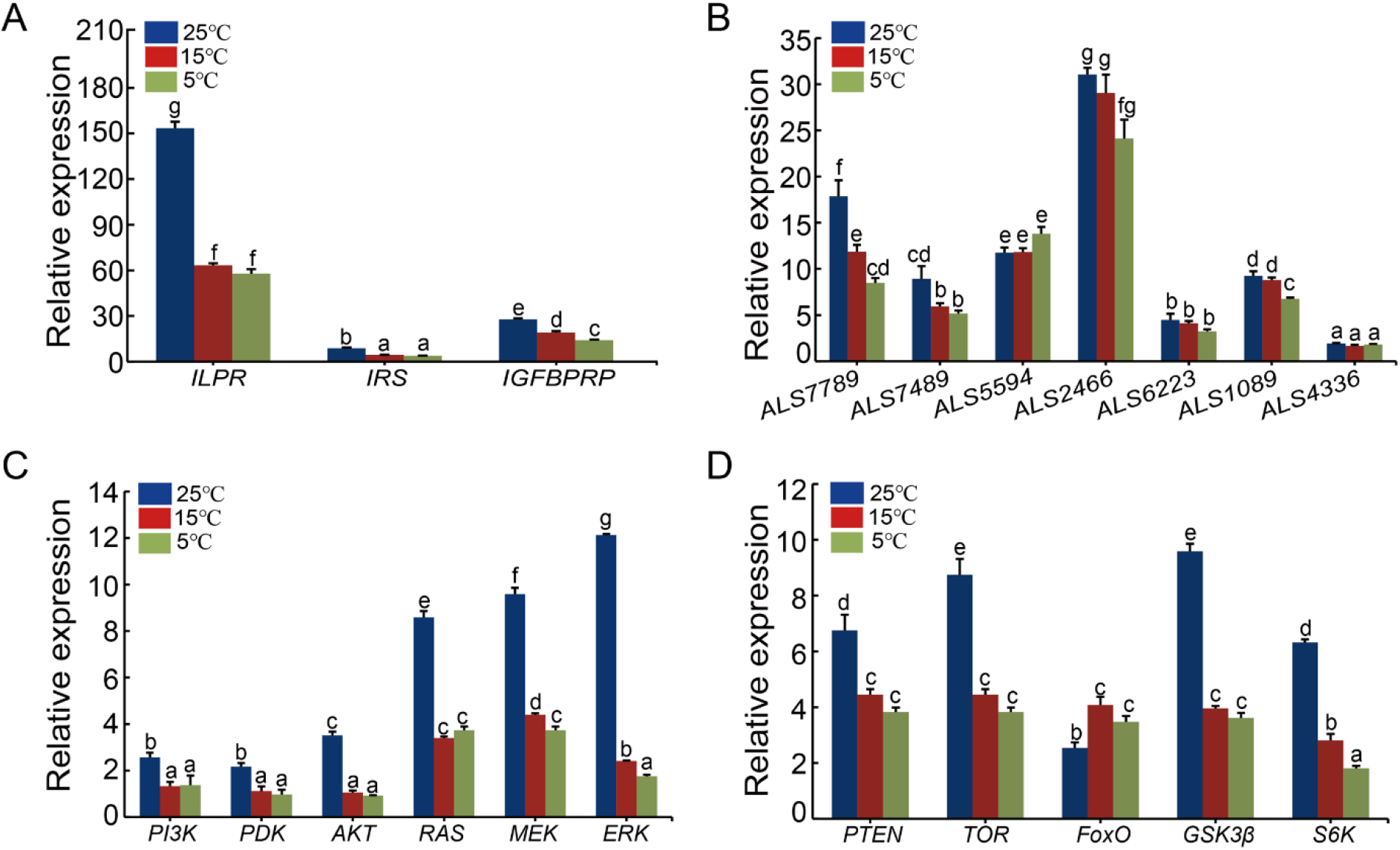
Effects of low temperature on the expressions of the IIS, PI3K-AKT, RAS-MAPK, and TOR signaling pathway genes. Data are expressed as the mean ± SD (n = 3). The significant difference (*P* < 0.05) among groups is indicated by the different lowercase letters.

## 4. Discussion

The involvement of insulin/insulin-like growth factor (IIS) signaling pathway in regulating growth and energy metabolism has been well-studied in vertebrates, while the role of IIS pathway in growth regulation of marine invertebrates such as oysters remains largely unexplored. In a previous study, we identified four insulin-like peptide genes in *C. gigas*, and showed that their expression levels were greatly associated with growth. While how the insulin-like peptides function to play roles in growth regulation of *C. gigas* deserves further investigation. In the present study, we performed an extensive bioinformatics data mining and identified one *ILPR*, one *IRS*, one *IGFBP*, and seven *IGFALSs* genes in *C. gigas*. Phylogenetic analysis confirmed their identities and reconstructed their evolutionary relationships, respectively. Expression profiling of the major genes of IIS and ILPR-mediated signaling pathways, including PI3K-AKT and RAS-MAPK signaling pathways, were all expressed at higher levels in the fast-growing “Haida No.1” oysters. Furthermore, expression levels of IIS, PI3K-AKT, RAS-MAPK and TOR signaling pathway genes were significantly affected by nutrient abundance and temperature in a gene-specific manner. These results suggested the critical roles of IIS signaling pathway in growth regulation of *C. gigas*, and confirmed the conserved role of the IIS and ILPR-mediated signaling pathways in growth control among various organisms.

Identification of IIS, PI3K-AKT, and RAS-MAPK signaling pathway genes in *C. gigas* suggested the conserved role of the IIS in growth and metabolism regulation. Phylogenetic analysis showed that the only one *C. gigas* ILPR was clustered with the ILPRs from other invertebrates and had a relatively closer homology with the insulin receptor (INSR) in comparison with other members of the IR subfamily in vertebrates, including the IGF1R, IGF2R and INSRR, indicating the similar physiological function between the ILPR in *C. gigas* and the insulin receptor in vertebrates. Notably, although four insulin-like peptide genes were found in *C. gigas*, there is only one ILPR, which was strikingly different from the case in vertebrates, deserving further investigations on the evolution of ligand and receptor binding for insulin-like peptides. The IRS, which could bind to the phosphorylated tyrosine residues of beta subunits of the ILPR, is involved in transducing the IIS signaling pathway within the cell [42, 43]. In our present study, different from the vertebrates which have three IRS, only one IRS was identified in *C. gigas*, indicating specific binding of IRS to beta subunits of ILPR in *C. gigas*.

The IGFBP family is composed of six distinct types of IGFBP (IGFBP1-6) and ten IGFBPRP in vertebrates, which is evolutionarily ancient and conserved [9]. In invertebrates, there is no definite evidence indicating the existence of traditional IGFBP1-6 which could bind to IGF-I and IGF-II with higher affinity than the related proteins (IGFBPRP1 to IGFBPRP10) [44]. The IGFBP7, also known as IGFBPRP1, is distinct from other low-affinity IGFBPRP in that it can strongly bind to insulin [23], suggesting that IGFBP7 is likely to have special biological functions from other IGFBPs. In our present study, one IGFBPRP and seven ALSs were found in *C. gigas,* and the IGFBPRP was clustered into one clade with the IGFBPRP from other mollusks and IGFBP7 from vertebrates. Our analysis suggested that the *C. gigas* IGFBPRP had a similar evolutionary origin and physiological function with the IGFBP7 in vertebrates, supporting that the IGFBPRP (IGFBP7) may be the most ancient gene member among the IGFBP subfamily. Furthermore, the number of ALS is only one or two in vertebrates, while there are usually five to seven IGFALSs in arthropods and mollusks. The obviously larger number of IGFALSs in invertebrates than in vertebrates indicated that some of the ALSs could have been lost during the evolution after divergence of invertebrates and vertebrates.

Higher expression levels of insulin-like peptide genes were associated with fast-growth in the *C. gigas* as revealed in our previous study. In the present work, expression profiles of the major genes in the IIS signaling pathway were further determined to confirm the systemic regulatory role of IIS signaling pathway in growth regulation of oysters. Our results showed that the major genes of the IIS signaling pathway, including *ILPR, IRS* and *IGFBPRP* were all expressed higher in the fast-growing “Haida No.1” than wild oysters. However, the *C. gigas* ALSs showed distinct expression profiles. The ALS has been reported to play an indispensable role in growth regulation of vertebrates [45, 46]. In our present study, the *ALS5594* gene was expressed higher in wild oysters, while the *ALS7789* and *ALS7489* genes were expressed at higher levels in the “Haida No.1” oysters, indicating that the ALS also played important but diverse roles in growth regulation of *C. gigas*.

In vertebrates, once the ternary complex “IGFs/insulins-IGFBP-ALS” binds to the receptor, the receptor is auto-phosphorylated and recruits the IRS, then the IRS would trigger different downstream signaling pathways such as the PI3K-AKT and RAS-MAPK signaling pathways [47–49]. The MAPK and its downstream response elements are involved in cell growth and proliferation [50]. The activation of PI3K led to production of PIP3, which in turn promoted the activation of downstream factors such as AKT, TOR and GSK3β to regulate cell proliferation, energy metabolism and hormone secretion [51, 52]. In our present study, we found that the ILPR mediated signaling pathway was conserved among vertebrates and invertebrates. The major genes of PI3K-AKT and RAS-MAPK signaling pathways, including the *PI3K*, *PDK*, *AKT*, *RAS*, *MEK*, *ERK*, *TOR*, and *GSK3β* were all expressed at higher levels in “Haida No.1” variety, while the negative regulators *PTEN, FoxO* and *S6K* were all highly expressed in wild oysters. Therefore, ILPR is speculated to transduce signaling from the active ternary complex “ILP-IGFBP-ALS” to the downstream PI3K-AKT and RAS-MAPK signaling pathways, functioning to regulate energy metabolism and cell proliferation in oysters (Fig. 6).

**Fig. 6.**
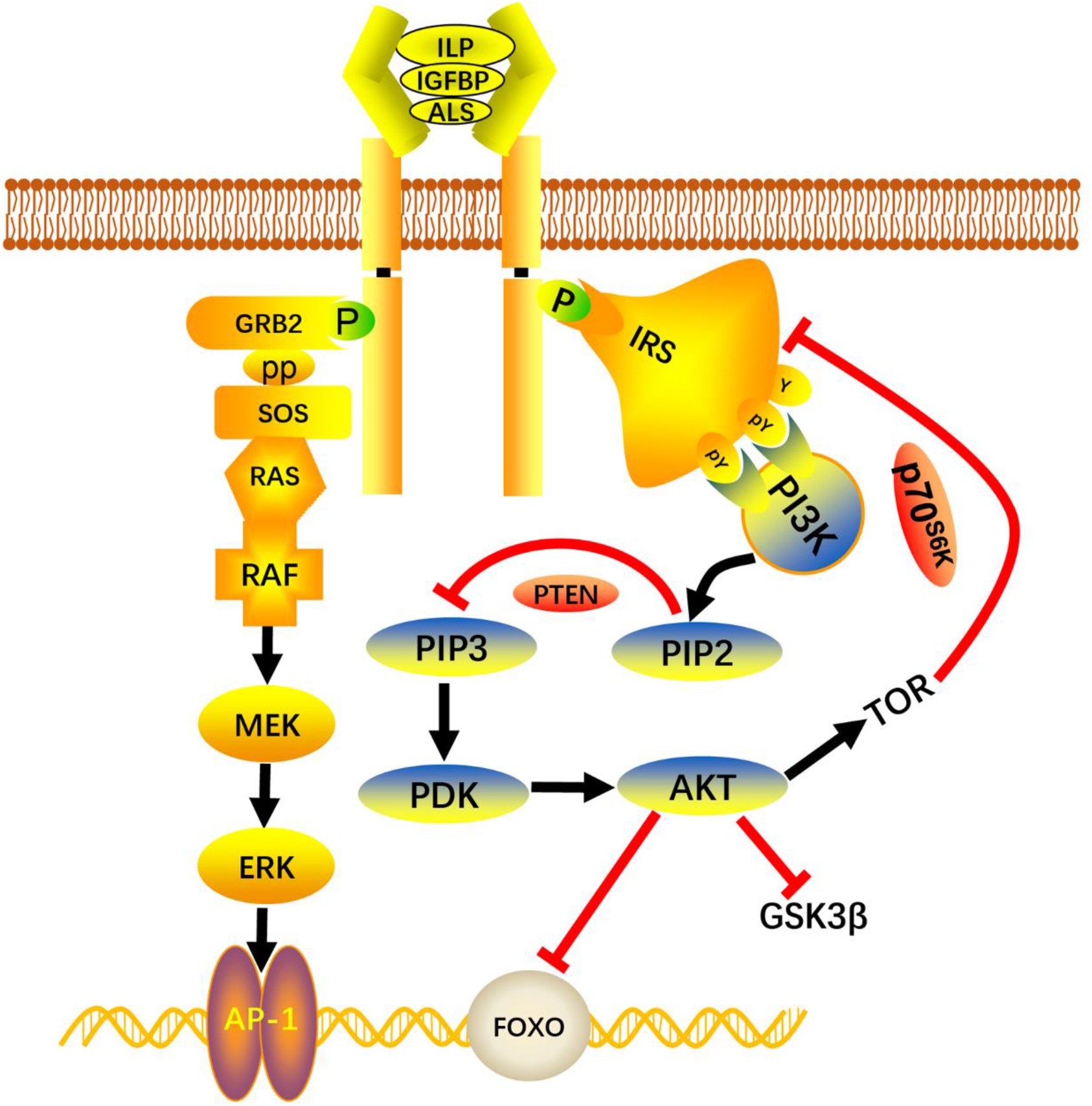
The proposed model of ILPR mediated IIS signaling pathway in growth regulation of *C. gigas*. The binding of insulin-like peptide to its receptor (ILPR) induces conformational changes in the alpha and beta subunits of the receptor. The IRS binds to the activated sites of the beta subunits and transduces signal to the cell through the PI3K-AKT and RAS-MAPK signaling pathways. The other elements including PTEN, GSK3β, FoxO, TOR could regulate the activation of the PI3K-AKT signaling pathway in *C. gigas* through a similar way as in vertebrates. The rapamycin-sensitive TORC1 (TOR complex 1) activates the translational regulator S6K (S6 kinase), leading to increased protein synthesis in the presence of nutrients, and the S6K could inhibit the activity of the IRS, finally regulate the PI3K-AKT and RAS-MAPK signaling pathways.

Fasting/re-feeding and low temperature culture experiments were performed to further confirm the speculation. In our present study, the major genes of IIS signaling pathway including the *ILPR*, *IRS*, *IGFBP*, and the seven *ALSs* all showed different expression patterns during fasting and re-feeding process. The expressions of *ILPR*, *IRS*, *IGFBP*, *ALS5594*, *ALS7489*, *ALS2466* and *ALS1089* genes all decreased once food was deprived and recovered to normal level once re-feeding, which were consistent with the expression profile of insulin-like peptide (ILP) gene in our previous study [40]. These results suggested that ILP, ILPR, IRS, IGFBP and ALSs may play systemic roles in food intake, nutrient digestion and absorption through the ternary complex “ILP-IGFBPRP-ALS” in *C. gigas*. In addition, the expressions of *ALS7789* and *ALS4336* genes were all up-regulated during fasting and re-feeding process, which were consistent with the expression profiles of *MIRP3, MIRP3-like* and *ILP7* genes in our previous study [40]. We speculate that the ALS7789 and ALS4336 may be associated with physiological role of molluscan insulin-related peptide 3 (MIRP3), molluscan insulin-related peptide 3-like (MIRP3-like) and insulin-like peptide 7 (ILP7), and are indispensable in energy metabolism under the control of neuroendocrine activity. Furthermore, low temperature greatly suppressed the expressions of *ILPR*, *IRS*, *IGFBP*, *ALS7789*, *ALS7489*, and *ALS1089* genes. These results suggested that the IIS signaling pathway is responsive to fluctuations of nutrition condition and ambient temperature.

The key elements of the PI3K-AKT and RAS-MAPK also showed different expression patterns during fasting and re-feeding process. The TOR nutrient pathway, which is regulated by insulin, nutrient abundance, energy and growth factors, function to modulate insulin-stimulated glucose transport, cellular metabolism, growth and proliferation [2]. In addition, the TORC1 activate the translational regulator S6K (S6 kinase), leading to increase protein synthesis in the presence of nutrients [53–56]. In our study, the expressions of *TOR* and *S6K* genes were all decreased during fasting and recovered to normal level once refeeding. GSK3β phosphorylates serine and threonine sites of a variety of substrates, including glycogen synthase, to regulate glycogen synthesis [58]. In our previous study, the expression of *GSK3β* was inhibited under food deprivation condition and recovered to normal level once the food was abundant, indicting the reduction of glycogen synthesis under food deprivation condition. As the negative regulator, FoxO regulates cell proliferation and energy metabolism through the transcriptional activation of certain genes [58]. In our study, the expression of *FoxO* was up-regulated during fasting and down-regulated once re-feeding. Furthermore, the expressions of IIS and ILPR mediated signaling pathway genes were all inhibited under low culture temperature except for the negative regulator *FoxO*. These results confirm that the IIS and ILPR mediated signaling pathways are conserved and ILPR could transduce signaling from the active ternary complex “ILP-IGFBP-ALS” to the downstream PI3K-AKT and RAS-MAPK signaling, functioning to regulate the energy metabolism and cell proliferation of *C. gigas* (Fig. 6).

In summary, we identified and characterized one *ILPR*, one *IRS*, one *IGFBPRP*, and seven *IGFALS*s in *C. gigas*. Expression profiling of these genes between fast-growing oysters and slow-growing wild oysters suggested their critical roles in growth regulation. The nutrient abundance and ambient temperature greatly affected the expression profiles of these genes. Furthermore, major genes of the ILPR-mediated PI3K-AKT and RAS-MAPK signaling pathways were all expressed at higher levels in the fast-growing “Haida No.1” variety, and were significantly suppressed under low temperature condition. These observations suggest that the IIS signaling pathway is involved in the growth regulation of *C. gigas* through regulating food intake, nutrition metabolism and cell proliferation. This work provid valuable information for further investigation on growth regulation mechanism in mollusks as well as other invertebrates.

## Acknowledgments

This work was supported by the grants from National Natural Science Foundation of China (Nos. 31802293, 41976098 and 31741122), the Young Talent Program of Ocean University of China (No. 201812013), Laboratory for Marine Fisheries Science and Food Production Processes, Qingdao National Laboratory for Marine Science and Technology (No. 2017-2A04), and China Postdoctoral Science Foundation (No. 2017M622283).

## Author contributions

SL conceived and designed the study. YL, HF, FZ, LR, and JT collected the samples and executed the experiments. YL, HF and SL analyzed the data. YL drafted the manuscript, and SL revised the manuscript. QL provided reagents and materials and supervised the study. All authors have read and approved the final version of the manuscript.

## Availability of data and materials

All data generated and analyzed during this study are included in this published article and its supplementary files.

