## Supplemental Tables and Figures for "Insulin-like peptide receptor mediated signaling pathways orchestrate regulation of energy homeostasis in the Pacific oyster, *Crassostrea gigas*"

**Supplementary Table 1.** The gene names and accessions of the sequences used in the phylogenetic analyses (Provided in a separate EXCEL file).

**Supplementary Table 2**. Primers used for the real-time PCR in this study.

| Gene name | Forward primer (5'-3') | Reverse primer (5'-3') |
|---|---|---|
| *ILPR* | ACCAGGGACTGTCCAATGAG | GGTCTGTAGCGCCAACATTT |
| *IRS* | TGTGGGTCAGGAAGGGAATC | CCAAACGAGCCTGACCTAGA |
| *IGFBPRP* | ACCTCGCCTGTAAGATGGAC | TCGGTACCACAGAGTGTGTC |
| *ALS5594* | GGCCGCTTATGCTAATGGAG | ACTCGTCCAACACATCCTGT |
| *ALS7789* | GTCTCCCGAGGGACATACTG | GTGCAACGTCTGAATCTCGT |
| *ALS7489* | TTCCTCGAGAACACGGACAT | AGGTGTTGGAAGGTGTAGGG |
| *ALS2466* | AACAGGCACTACCCAAGGTT | TAGAGGAAGCGGTGGTTTGT |
| *ALS6223* | CGAGCGTCAAAGATGTCCTG | TCGATCCCGATCCGTTTGAT |
| *ALS1089* | TCAACAACACAGCGTGCTTA | CATTTGAAAGCGGTGCCATC |
| *ALS4336* | CCACAGAACGCCTTTGAGAG | GGGAACTGCTGAACGAACTC |
| *PI3K* | TCCGTTCACATACCAAACGC | GGGCAGAGTGGCTTCATAGA |
| *PDK* | CATCAAGGTGCTGGAGAAGC | CTGTCTGTGTCCTGGAAGGT |
| *AKT* | AGAGGTTGCTCCAATCGTCA | ATGAAACACGCCATCAGCTC |
| *GSK3β* | CTAGCCTACATCCACTCGCA | AGGAGCCCTGTAGTAACGTG |
| *mTOR* | ACGTGACAAGACCTCCACAT | CAGGATCCCGGGAAGGTATC |
| *FoxO* | TCGCTACTACAGCGTTGTCT | GCCTGTCGTAGAAGCGAATC |
| *PTEN* | ACAAGATGGCGGATGTTGTG | GGGTTGTGGTCATCGAATGG |
| *RAS* | GACCCGACCATAGAGGACAG | TCCCTGCCCATTCTTCATGT |
| *MAPK* | TGGCTGTATCCTGGCAGAAA | TGTTCCATGGGACTTTGGGT |
| *ERK* | GCAATGGATCCACCACTTCC | ACGTGTTACCATGCACACTG |
| *EF1α* | AGTCACCAAGGCTGCACAGAAAG | TCCGACGTATTTCTTTGCGATGT |

**Fig. S1.** Phylogenetic analyses of the insulin receptor substrates.

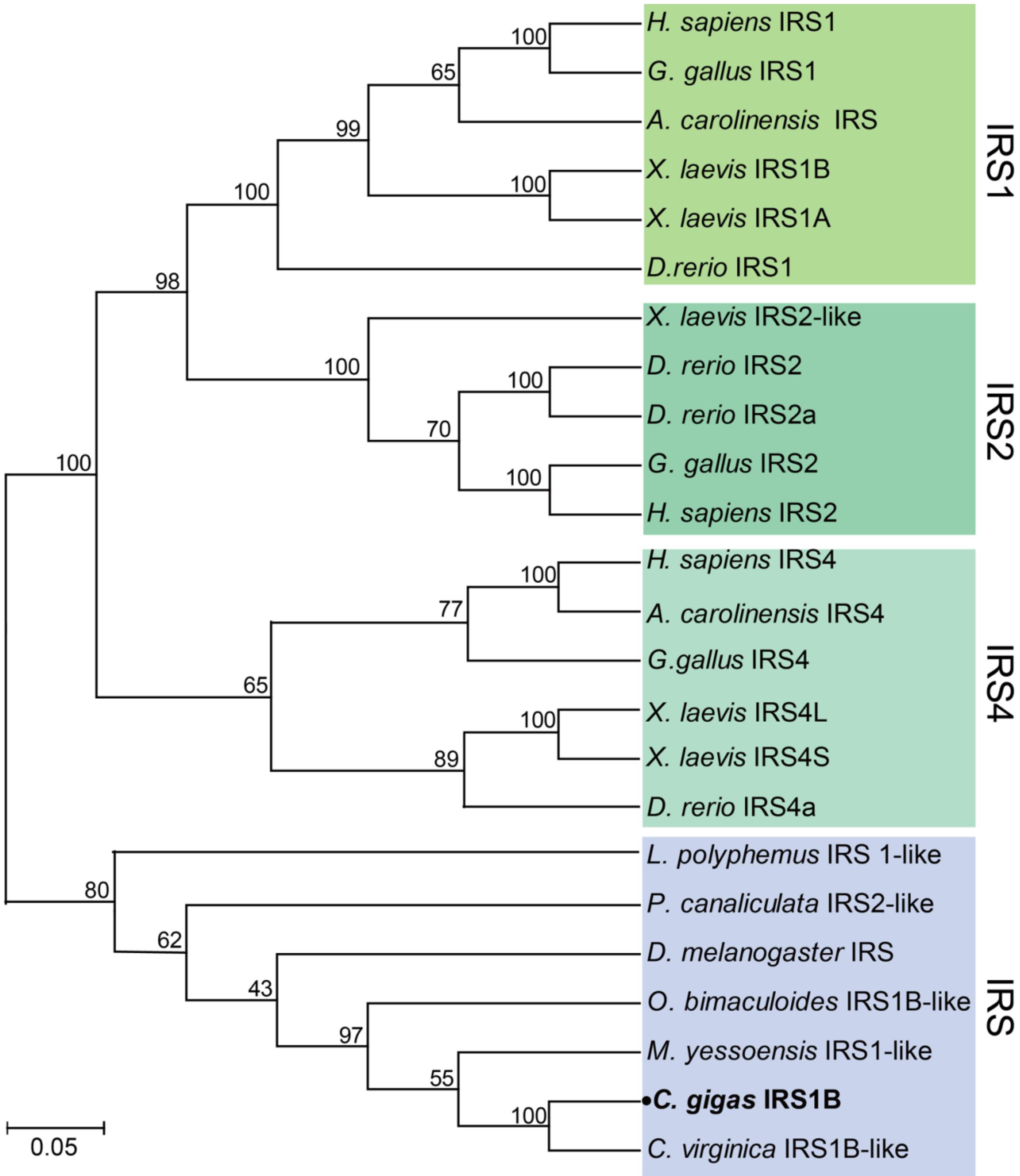

**Fig. S2.** Phylogenetic analyses of the IGFBP superfamily.

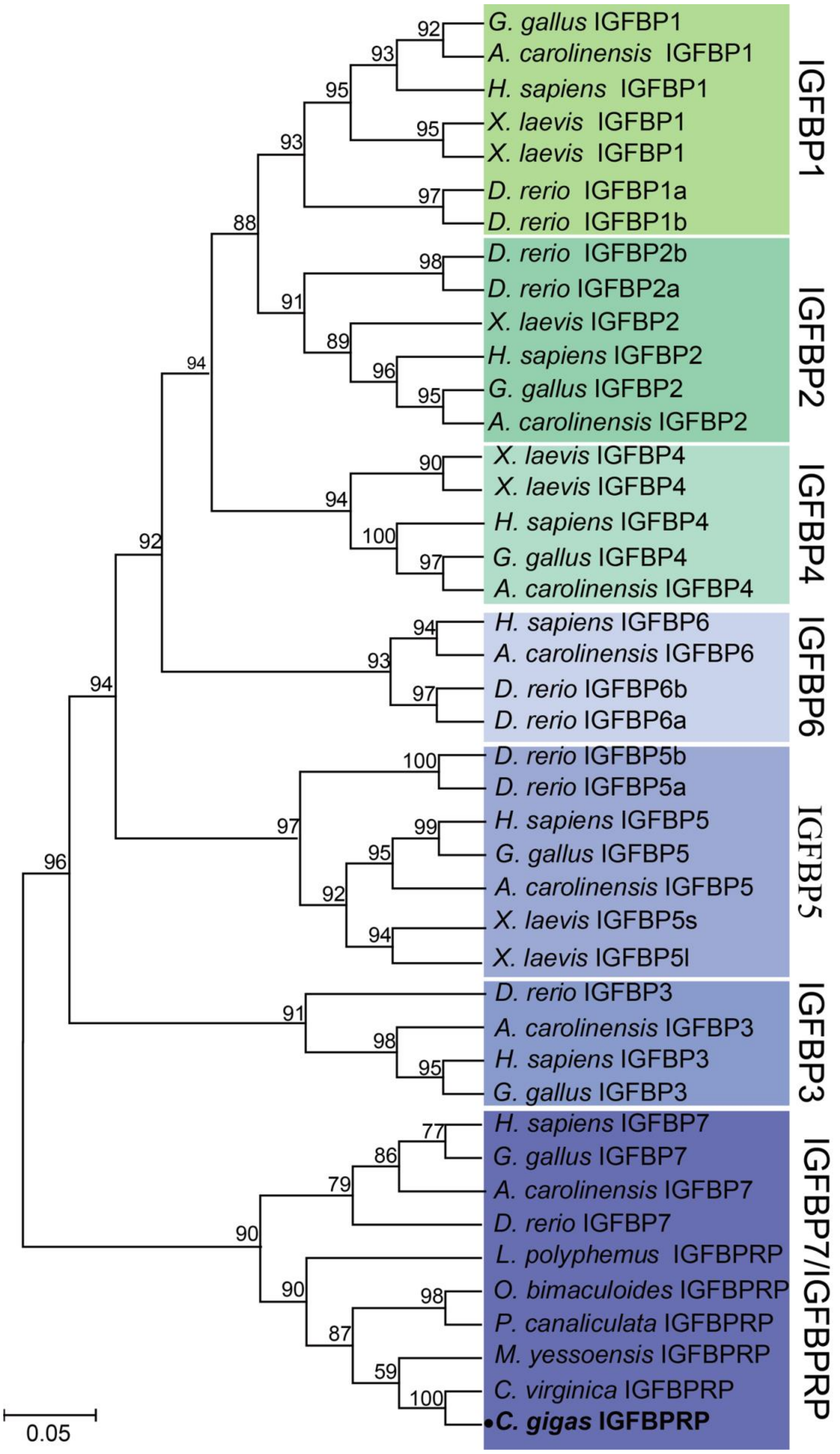

**Fig. S3.** Phylogenetic analyses of the IGF ALSs.

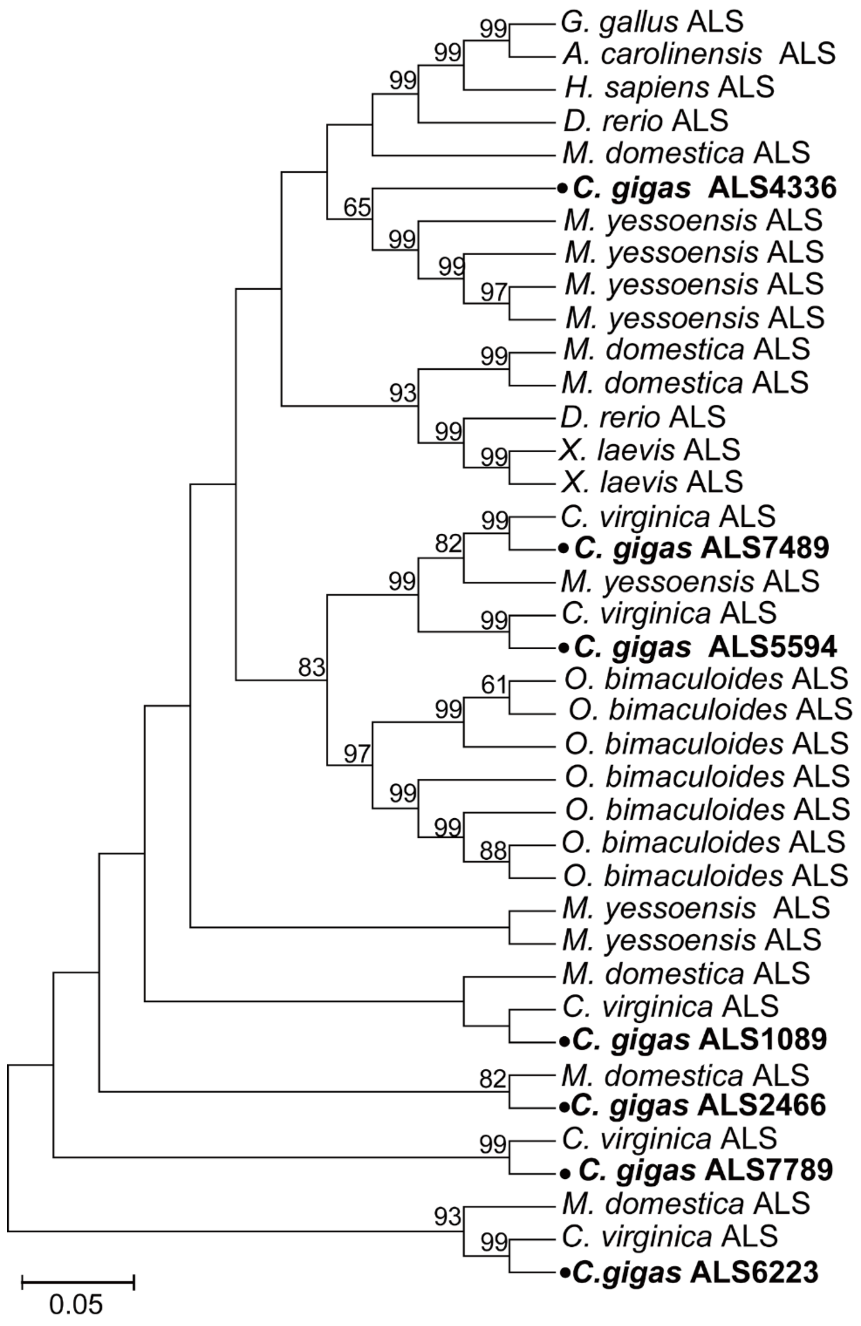
